# Treatment of the X chromosome in mapping multiple quantitative trait loci

**DOI:** 10.1101/2020.09.11.294017

**Authors:** Quoc Tran, Karl W. Broman

**Author notes:** **Corresponding Author:** Karl W Broman, Department of Biostatistics and Medical Informatics, University of Wisconsin–Madison, 2126 Genetics-Biotechnology Center, 425 Henry Mall, Madison, WI 53706.

## Abstract

Statistical methods to map quantitative trait loci (QTL) often neglect the X chromosome and may focus exclusively on autosomal loci. But the X chromosome often requires special treatment: sex and cross-direction covariates may need to be included to avoid spurious evidence of linkage, and the X chromosome may require a separate significance threshold. In multiple-QTL analyses, including the consideration of epistatic interactions, the X chromosome also requires special care and consideration. We extend a penalized likelihood method for multiple-QTL model selection, to appropriately handle the X chromosome. We examine its performance in simulation and by application to a large eQTL data set. The method has been implemented in the package R/qtl.

## INTRODUCTION

The X chromosome is often neglected in methods to map quantitative trait loci (QTL), yet it often requires special treatment. For example, in an intercross between two inbred strains, A and B, the offspring have genotypes AA, AB, or BB on the autosomes, but on the X chromosome males are hemizygous A or B, while females have genotypes either AA or AB, if their paternal grandmother was from strain A, or genotypes AB or BB, if their paternal grandmother was from strain B. These differences introduce three difficulties: the treatment of the male hemizygous genotypes, the potential for spurious evidence for X chromosome linkage due to a sex or cross-direction difference in the phenotype, and the need for separate thresholds for statistical significance for the X chromosome and the autosomes, due to a difference in degrees of freedom in the linkage tests. Similar considerations apply in genome-wide association studies (Zheng *et al.* 2007; Clayton 2008; Hickey and Bahlo 2011).

Broman *et al.* (2006) described the necessary modifications to single-QTL analysis by interval mapping (Lander and Botstein 1989). They identified additional covariates that need to be considered in the null model with no QTL, to avoid spurious linkage. They identified appropriate linkage tests for various configurations of an intercross or backcross, ensuring that the null model is nested within the alternative, single-QTL model. And they described an approach to obtain separate permutation-based significance thresholds for the autosomes and X chromosome.

Many of the difficulties with the X chromosome are further exacerbated in the consideration of multiple-QTL models. Multiple-QTL models have a number of advantages for QTL mapping, including the potential for increased power to detect QTL, the ability to separate linked QTL, and the possibility of identifying epistatic interactions among QTL. But, for example, in the consideration of epistatic interactions, the degrees of freedom for the test for epistasis can vary dramatically depending on whether both loci are on autosomes, both are on the X chromosome, or one is on the autosome and one is on the X chromosome (Broman and Sen 2009).

We describe an approach for multiple-QTL model selection, including the investigation of epistatic interactions, that deals appropriately with the X chromosome. We build upon the penalized likelihood approach of Broman and Speed (2002) and Manichaikul *et al.* (2009). We use permutation tests (Churchill and Doerge 1994) with two-dimensional, two-QTL genome scans, to derive separate thresholds for the main effects for QTL on the autosomes and X chromosome, and for pairwise epistatic interactions, depending on whether both, one, or neither QTL is on the X chromosome.

We use computer simulations to assess the performance of our approach. We further illustrate the approach through application to a large expression quantitative trait locus (eQTL) study. The method has been implemented in the widely-used package R/qtl (Broman *et al.* 2003).

## METHODS

We consider the case of a backcross or an intercross derived from two inbred lines and of a continuously varying quantitative trait with normally distributed residual variation. We focus on Haley-Knott regression (Haley and Knott 1992), which handles missing genotype information by considering a regression of the phenotype on QTL genotype probabilities, calculated conditional on the available marker genotypes (see Broman and Sen 2009). Thus we consider models of the form *y* = *Xβ* + *ϵ* where the covariate matrix *X* includes an intercept and, in a backcross, includes one column for each autosomal QTL, while in an intercross, it includes two columns for each autosomal QTL. For X chromosome QTL, we allow for QTL *×* sex interactions, for example allowing that male hemizygotes may have different average phenotypes than female homozygotes. In an intercross with both sexes and both cross directions, we would include sex and cross-direction covariates (see Broman *et al.* 2006), plus two columns of 0/1 indicators for the female genotypes and one such column for the male genotypes. For models with epistatic interactions, we impose a hierarchy on the models, with the inclusion of a pairwise interaction requiring the inclusion of both corresponding main effects.

Broman and Speed (2002) introduced the use of a penalized LOD score criterion for multiple-QTL model selection in this context. They focused on additive QTL models and placed a linear penalty on the number of QTL. Their criterion was

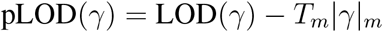

where *γ* denotes an additive QTL model, *|γ|*_*m*_ is the number of QTL in the model, and LOD(*γ*) is the log_10_ likelihood ratio for the model *γ* versus the null model of no QTL. The penalty *T*_*m*_ was chosen as the 1 *− α* quantile of the genome-wide maximum LOD score in a permutation test (see Churchill and Doerge 1994), and they sought the model with maximum pLOD.

This approach could be modified to allow different penalties for QTL on the X chromosome than those on autosomes, using significance thresholds as in Broman *et al.* (2006), with *T*_*mA*_ and *T*_*mX*_ being the 1 *− α*_*A*_ and 1 *− α*_*X*_ quantiles, respectively, of the maximum LOD scores across the autosomes and the X chromosome, from a permutation test. Broman *et al.* (2006) took 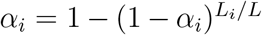, where *L*_*X*_ is the length of the X chromosome, *L*_*A*_ is the total lengths of the autosomes, and *L* = *L*_*A*_ + *L*_*X*_.

Manichaikul *et al.* (2009) extended the penalized LOD score approach to consider models with pairwise interactions among QTL. They imposed a hierarchy on the models, with the inclusion of a pairwise interaction requiring the inclusion of both corresponding main effects. We will also impose this hierarchy. They used the same main-effect penalty as in Broman and Speed (2002), but added a penalty on interaction terms. They considered heavy and light penalties on interactions. The heavy interaction penalty was taken as the 1 *− α* quantile of the LOD score for the interaction term in a two-dimensional, two-QTL scan of the genome, under the null hypothesis of no QTL. The light interaction penalty was derived as the 1 *− α* quantile for the LOD score comparing a two-locus interactive model to a single-QTL model, but then subtracting off the main effect penalty.

The light interaction penalty has the advantage of giving greater power to detect interactive QTL, but with an increased rate of false interactions. Exclusive use of the light penalty gave a high rate of false positive QTL, and so they used a compromise: considering a QTL model as a graph with nodes being QTL and edges being pairwise interactions, they allowed no more than one light interaction penalty for each connected component and placed heavy penalties on all other interactions.

In adapting the approach of Manichaikul *et al.* (2009) to handle the X chromosome, we propose a similar penalized LOD score, but with separate thresholds for interactions in three regions A:A, A:X, X:X. Broman *et al.* (2006) identified sex and cross-direction covariates that should be included under the null hypothesis; we include these covariates under all models. The ad hoc system of heavy and light penalties on interaction terms, suggested by Manichaikul *et al.* (2009), becomes unmanageable when separate interaction penalties are considered for the three regions, and so we allow light penalties only for interactions for which both QTL are on autosomes; any QTL involving the X chromosome must be a heavy penalty.

The main effect penalties are as described above. The interaction penalties are based on significance levels *α*_*AA*_, *α*_*AX*_, *α*_*XX*_, based on the areas of the corresponding regions. For region *i*, we use 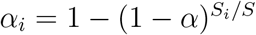, where 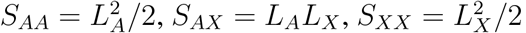, and *S* = *S*_*AA*_ + *S*_*AX*_ + *S*_*XX*_.

In the following, we denote the method of Manichaikul *et al.* (2009), treating the autosomes and X chromosome the same, as XeqA (for X chromosome and autosomes treated equally), and the proposed approach, treating the X chromosome separately, as XneA.

The proposed approach requires a permutation test with a two-dimensional, two-QTL genome scan, keeping track of the maximum LOD score in the three regions, A:A, A:X, and X:X. And as the quantiles for the A:X and X:X regions are much farther out in the tail, we will need to perform many more permutations for the A:X and X:X regions in order to get accurate estimates of the penalties. In Broman *et al.* (2006), they suggested using a factor *L*_*A*_/*L*_*X*_ additional permutations for the X chromosome as for autosomes. Following that approach, we would require (*L*_*A*_/*L*_*X*_)^2^ as many permutations for the X:X region as for the A:A region.

To search the space of QTL models, we use forward selection up to a model with 10 QTL, followed by backward elimination to the null model. At each step of forward selection, we scan the genome for an additional additive QTL, and then for each QTL in the current model, we scan the genome for a QTL that interacts with that QTL. We step to the model that gives the largest penalized LOD score, even if it results in a decrease relative to the current model. In the end, we select the model with maximum penalized LOD score, among all models considered in the search.

### Data and software availability

Software implementing the proposed methods have been incorporated into the R/qtl package, which is available at its website (https://rqtl.org), at GitHub (https://github.com/kbroman/qtl), and at the Comprehensive R Archive Network (CRAN, https://cran.r-project.org). The data from Tian *et al.* (2016), considered in the Application section below, are available at the Mouse Phenome Database, at https://phenome.jax.org/projects/Attie1. The detailed code we used for the simulations and in the application are available at https://github.com/kbroman/Paper_qtlX.

## SIMULATIONS

To investigation the performance of our proposed approach, we conducted a computer simulation. We compare the Type I error rate and power, including the relative rates of Type I errors on autosomes versus the X chromosome.

We consider a backcross with 250 individuals, split evenly between males and females, using a genome modeled after the mouse, with markers at a 10 cM spacing. We considered a variety of small QTL models: a single QTL, two additive QTL, or three additive QTL all on autosomes; a single QTL or two additive QTL on the X chromosome; two interacting QTL on autosomes; and two interacting QTL with one on an autosome and one on the X chromosome. In all cases, the percent phenotypic variance explained by each QTL was approximately 8%. We used Haley-Knott regression (Haley and Knott 1992) and stepwise model selection. Analyses were performed with R/qtl version 1.42-8 (Broman *et al.* 2003) and R version 3.5.3 (R Core Team 2020).

First we use simulation to derive the various penalties for the XeqA and XneA methods. For the XneA methods, we performed 1,056 simulation replicates for the A:A region, 8,544 replicates for the A:X region, and 276,192 replicates for the X:X region. The estimated penalties are shown in Table 1. These penalties are used to perform stepwise model selection. The results, based on 1,000 simulations, are displayed in Figure 1. Figure 1A shows the Type I error rate for main effect QTL. Treatment of the X chromosome separately from the autosomes (XneA) results in a smaller Type I error rate than when all chromosomes are treated equally (XeqA). The XeqA method has high Type I error rate particularly when the simulated model has two main QTL on X chromosome.

**Table 1:** Estimated penalties for the simulations and the application, for the XeqA and XneA methods.

| **Penalty** | **Simulations** |  | **Application** |  |
| --- | --- | --- | --- | --- |
|  | **XeqA** | **XneA** | **XeqA** | **XneA** |
| $T_{mA}$ | 2.98 | 2.88 | 3.85 | 3.90 |
| $T_{mX}$ | | 3.61 | | 3.51 |
| $T_{iAA}^H$ | 4.83 | 4.34 | 6.69 | 6.59 |
| $T_{iAA}^L$ | 2.28 | 1.83 | 3.73 | 3.62 |
| $T_{iAX}$ | | 5.74 | | 7.21 |
| $T_{iXX}$ | | 5.15 | | 4.88 |

**Figure 1:**
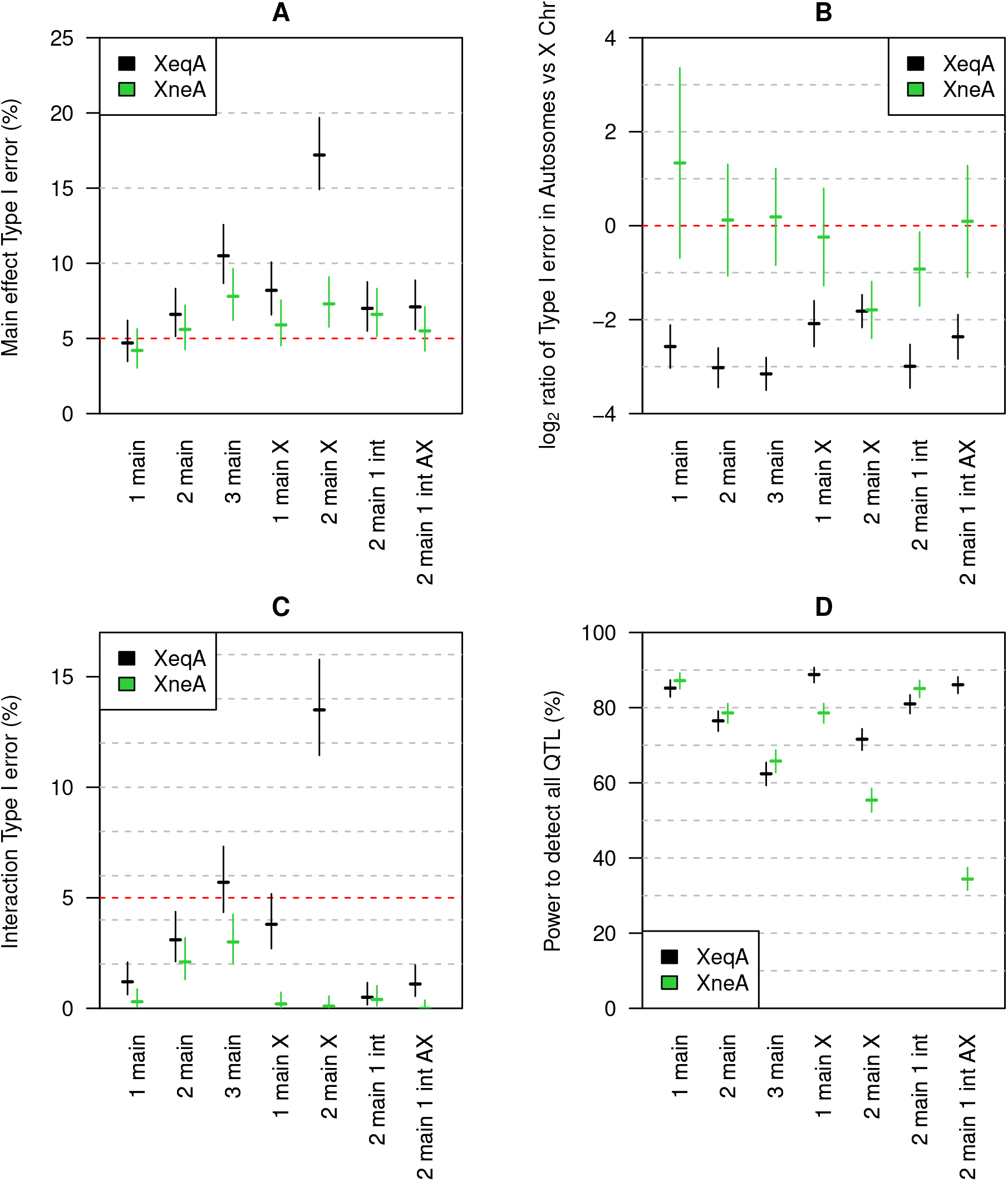
Simulation results: Type I error and power of these methods across simulated models. Black is the method treating the X chromosome and autosomes the same (XeqA); green is the method treating the X chromosome and autosomes differently (XneA). The estimates and 95% confidence intervals are based on 1000 simulations. A: Type I error for main effects. B: log_2_ ratio of the Type I error of main effects for autosomes versus the X chromosome. C: Type I error for interactions. D: Power to detect QTL.

Figure 1B shows the log_2_ ratio of Type I error rates, for the autosomes versus the X chromosome. The proposed XneA method serves to balance type I error rates on the autosomes and the X chromosome, while the XeqA method shows increased type I error on the X chromosome.

Figure 1C shows the type I error rate for interactions. The XneA method is conservative, with below-target rates of type I errors. The XeqA method has high false type I error rates for interactions, particularly in the case of two QTL on the X chromosome.

Figure 1D shows the power to detect QTL. For autosomal QTL, the two approaches have similar power. The XneA method has lower power to detect X chromosome QTL. This is the trade-off to control the Type I error rate on the X chromosome.

## APPLICATION

To illustrate our method regarding the X chromosome in multiple-QTL model selection, we consider the eQTL data of Tian *et al.* (2016). The data concern a mouse intercross between the diabetes-resistant strain C57BL/6J (abbreviated B6) and the diabetes-susceptible strain BTBR *T* ^+^*tf* /J (abbreviated BTBR). There were approximately 500 F_2_ offspring, all genetically obese through introgression of the leptin mutation (*Lep*^*ob/ob*^). Genome-wide gene expression data were available on six tissues, assayed with custom two-color Agilent microarrays; here, we focus on liver expression, for which there was data on 483 F_2_ mice. Mice were genotyped with the Affymetrix 5K GeneChip, giving 2,057 informative markers.

The F_1_ parents were derived from a cross between a female BTBR and a male B6. Thus the female F_2_ offspring all have an X chromosome from BTBR and so are either homozygous BTBR or heterozygous. The males have a Y chromosome from B6 and are hemizygous B6 or BTBR on the X.

We first used computer simulation to estimate penalties for the XeqA and XneA methods. For all mice in the study, we simulated standard normal phenotypes, independent of genotype, and performed two-dimensional, two-QTL genome scans with sex as an additive covariate to estimate the various penalties. We performed 1,056 simulation replicates for the A:A region, 9,504 replicates for the A:X region, and 335,328 replicates for the X:X region. The numbers of replicates were chosen based on the areas of the corresponding regions, with *L*_*A*_/*L*_*X*_ ≈ 19. Penalties using *α* = 0.05 are displayed in Table 1. The main effect penalty on the X chromosome is lower than that on the autosomes, because of reduced recombination. (Each F_2_ mouse has a single recombinant X chromosome.) The interaction penalty for two QTL on the X chromosome is lower than the heavy interaction penalty for two QTL on autosomes, but the penalty for interactions between autosomal and X chromosome QTL is highest.

We considered the 37,827 gene expression microarray probes with known genomic location (including 1,433 on the X chromosome, 21 on the Y chromosome, and 9 mitochondrial probes). The gene expression values were transformed to normal quantiles. We performed single-QTL genome scans and identified 10,814 microarray probes (29%) with at least one significant QTL at *α* = 0.05, allowing separate significance thresholds on the autosomes and X chromosome. For the multiple-QTL analyses, we will focus on these 10,814 microarray probes.

Table 2 shows the distribution of the estimated number of QTL by the two methods, as well as the estimated number of pairwise interactions. As we focused on microarray probes with at least one significant QTL in a single-QTL genome scan allowing separate autosome and X chromosome thresholds, it is no surprise that all of these microarray probes showed at least one QTL by the XneA method.

**Table 2:** Distribution of the estimated number of QTL and pairwise interactions for the liver expression data from Tian *et al.* (2016), for the 10,814 gene expression microarray probes with at least one QTL by the XneA method.

| **No. QTL** | **XeqA** | **XneA** | **No. interactions** | **XeqA** | **XneA** |
| --- | --- | --- | --- | --- | --- |
| 0 | 56 | 0 | 0 | 10408 | 10419 |
| 1 | 5212 | 5432 | 1 | 392 | 380 |
| 2 | 2566 | 2518 | 2 | 12 | 13 |
| 3 | 1444 | 1397 | 3 | 2 | 2 |
| 4 | 732 | 712 |  |  |  |
| 5 | 390 | 373 |  |  |  |
| 6 | 205 | 194 |  |  |  |
| 7 | 114 | 116 |  |  |  |
| 8 | 46 | 45 |  |  |  |
| 9 | 26 | 25 |  |  |  |
| 10 | 23 | 2 |  |  |  |

There were 56 microarray probes that showed no QTL by the XeqA method. All of these showed a single QTL on the X chromosome by the XneA method. This can be explained by the fact that the main effect penalty for the X chromosome is considerably smaller than that for the autosomes (Table 1).

In general, these expression traits show a large number of QTL. While about half show a single QTL, about a quarter show 2 QTL and another quarter show 3 or more QTL. The number of interactions, however, is quite small. Less than 4% of expression traits show an interaction, and about 0.1% have two or more interactions.

The two methods gave identical results for the vast majority of the microarray probes: 10,290 (95.2%) gave the same QTL model by the two methods, and just 524 (4.8%) gave different models. For 257 microarray probes (2.4%), the XneA results were nested within the XeqA results, while for 98 microarray probes (0.9%), the XeqA results were nested within the XneA results. Another 169 (1.6%) had more complex differences, with some gains and losses on both sides. In 139 of the 524 cases where the two methods gave different results, part of the difference included the X chromosome: 86 microarray probes gained an X chromosome QTL by the XneA method, while 53 lost an X chromosome QTL.

As one example of the kinds of differences observed, consider microarray probe 10003837305, which is on chromosome 1 at 152 Mb (63.2 cM), though not in a known gene. The results by the two methods are in Table 3, with the results by the XeqA method on the left, and those for the XneA method on the right. By the XeqA method, there are two QTL on chromosome 7, one of which interacts with a locus on the X chromosome. With the XneA method, the QTL on chromosome 7 with the 7:X interaction is omitted, and instead there is an interaction between the other chromosome 7 QTL and the locus on chromosome 10. The position of the chromosome 10 QTL shifts a bit, but the positions of most other QTL do not change much. Note that the QTL on chromosome 1 is very close to the genomic location for this microarray probe.

**Table 3:** Comparison of the selected QTL models by the XeqA and XneA methods, on probe 10003837305.

| (a) XeqA |  |  |  |  | (b) XneA |  |  |  |  |
| --- | --- | --- | --- | --- | --- | --- | --- | --- | --- |
|  | name | chr | pos | LOD |  | name | chr | pos | LOD |
| Q1 | 1@60.0 | 1 | 60.0 | 5.4 | Q1 | 1@59.3 | 1 | 59.3 | 5.0 |
| Q2 | 7@2.5 | 7 | 2.5 | 10.7 | Q2 | 7@18.6 | 7 | 18.6 | 15.1 |
| Q3 | 7@20.1 | 7 | 20.1 | 5.8 | Q3 | 9@31.6 | 9 | 31.6 | 5.6 |
| Q4 | 9@31.6 | 9 | 31.6 | 7.1 | Q4 | 10@24.0 | 10 | 24.0 | 8.7 |
| Q5 | 10@3.0 | 10 | 3.0 | 4.0 | Q5 | 13@14.4 | 13 | 14.4 | 4.4 |
| Q6 | 13@14.4 | 13 | 14.4 | 5.4 | Q6 | 17@13.1 | 17 | 13.1 | 5.7 |
| Q7 | 17@19.6 | 17 | 19.6 | 7.1 | Q7 | X@56.4 | X | 56.4 | 34.5 |
| Q8 | X@57.4 | X | 57.4 | 42.1 | Q2:Q4 | 7@18.6:10@24.0 | 7:10 |  | 7.0 |
| Q2:Q8 | 7@2.5:X@57.4 | 7:X |  | 6.4 |  |  |  |  |  |

Figure 2 shows LOD profiles for the two models for this microarray probe, with results for the XneA method in green. Each curve shows the log_10_ likelihood when varying the position of the given QTL, keeping all other QTL fixed at their estimated positions, versus the model where the given QTL is omitted (along with any interactions it is involved in). These curves show the strength of evidence for the QTL as well as an indication of their mapping precision. Again, in switching from the XeqA method to the XneA method, one of the chromosome 7 QTL is lost, as is the 7:X interaction, but a 7:10 interaction is gained, and the position of the chromosome 10 QTL shifts.

**Figure 2:**
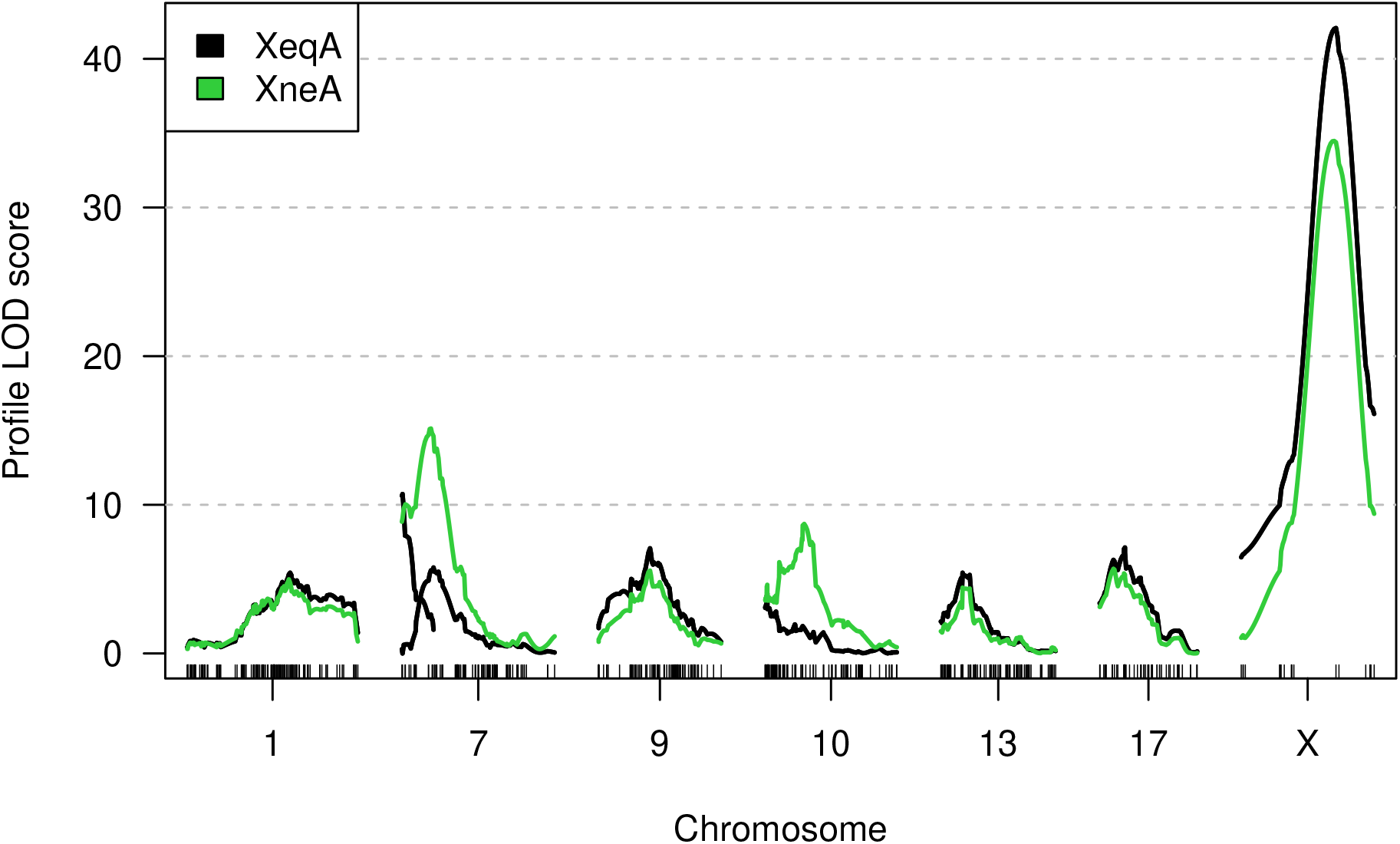
LOD profile of output models from XeqA (black) and XneA (green) methods on probe 10003837305. Note that there are two black curves on chromosome 7, because the XeqA method identified two QTL on that chromosome.

## DISCUSSION

We have introduced an approach for multiple-QTL model selection that takes account of the X chromosome. It builds on the penalized LOD score method of Manichaikul *et al.* (2009), but with a separate main-effect penalty for the autosomes and the X chromosome, and separate interaction penalties for the three kinds of pairwise interactions: between two autosomal QTL, between two X chromosome QTL, and between a QTL on an autosome and one on the X chromosome.

Our approach balances false positive rates between the autosomes and the X chromosome. In our computer simulations, which concerned a backcross with both sexes, the main effect penalty is higher for the X chromosome than for autosomes, and so the new approach gives a loss of power to detect QTL on the X chromosome. The application to the eQTL dataset of Tian *et al.* (2016) concerned an intercross with both sexes but just one cross direction, and so the main effect penalty was lower for the X chromosome than for autosomes, and so the new approach gave some additional QTL on the X chromosome.

In the eQTL application, for the vast majority of expression traits, the separate treatment of the X chromosome gave identical results. This is as expected: the penalties (Table 1) have relatively small changes, and these changes will only affect the results when loci or interactions have LOD scores in the small range between these penalty values.

In the eQTL application, many expression traits showed multiple QTL, including up to 10 QTL. (Our model search did not consider models with more than 10 QTL.) However, there were few expression traits for which we identified epistatic interactions. The high degrees of freedom of the test for epistatic interactions in an intercross, as well as large search space, result in large penalties on interactions and so low power to detect interactions.

Software implementing our approach has been incorporated into R/qtl (Broman *et al.* 2003), an add-on package to the general statistical software R (R Core Team 2020).

## ACKNOWLEDGMENTS

This work was supported in part by National Institutes of Health grants R01GM074244 and R01GM070683 (to K.W.B.). The authors thank two anonymous reviewers for comments to improve the manuscript.

